# A theoretical framework linking the effective number of breeders to demographic stochasticity in iteroparous species

**DOI:** 10.1101/2024.06.05.597532

**Authors:** Tetsuya Akita

**Author notes:** Corresponding author: Fisheries Resources Institute, Japan Fisheries Research and Education Agency, Kanazawa, 2-12-4, Japan. This manuscript was typeset using a slightly modified version of the LaTeX class file originally provided by Genetics Society of America. Journal branding has been removed for preprint dissemination purposes. This version does not imply acceptance or endorsement by the journal.

## Abstract

The effective number of breeders (*N*_b_) is widely used as a genetic summary statistic, yet its role in shaping demographic variability has remained underexplored. Here, we present a theoretical framework for iteroparous species that integrates *N*_b_ into the stochastic recruitment process of the number of adults (*N*) by modeling individual-level relationships between adult females and their offspring. We partition recruitment variation into parental, non-parental, and environmental components, and show that demographic stochasticity arising from non-Poisson reproduction, summarized by *N*_b_, can amplify environmental variance. Assuming an equal sex ratio and no sex-specific variation, we derive an approximation linking *N*_b_ to this amplification when *N*_b_/*N* < 0.1. For instance, *N*_b_ < 42 can increase recruitment variance by more than 5%. A key advantage of this approach is that *N*_b_ can be estimated from genetic data collected from a single cohort, making it applicable in data-limited or conservation-priority systems. Our framework highlights a distinct causal pathway by which increased reproductive variance contributes to demographic variability and extinction risk, especially in species where both *N*_b_ and *N*_b_/*N* are small. These findings provide new theoretical justification for using *N*_b_ as a life-history-independent metric to quantify demographic stochasticity and connect genetic monitoring with population viability analysis.

## Introduction

Effective population size (*N*_e_) determines the rate of genetic drift per generation (e.g. Charlesworth 2009; Waples 2025) and is widely used as an indicator of genetic diversity and reflects evolutionary vulnerability, particularly in conservation biology and wildlife management (Frankham *et al*. 2014). For example, the 50/500 rule concerning *N*_e_ (Franklin 1980) is one of the most well-known concepts, providing a widely accepted threshold for maintaining genetic health in the short and long term (Mace *et al*. 2008). Furthermore, since the adoption of the Convention of Biological Diversity Kunming-Montreal Global Biodiversity Framework (CBD 2022), which sets targets for conserving the genetic diversity of all species, the importance of *N*_e_ has received increasing attention in recent years. Although *N*_e_ has been proposed as a proxy for census size or adult number (*N*) (e.g. a conservative estimate of *N*_e_/*N* ≈ 0.1 for conservation assessments of genetic diversity, Hoban *et al*. 2020, 2024), theoretical studies that link *N*_e_ to population persistence independently of fitness-related effects remain scarce.

Furthermore, *N*_e_/*N* serves as an indicator of the variance in reproductive success (Wang *et al*. 2016; Waples 2016). For instance, many marine organisms with high fecundity and mortality exhibit extremely low *N*_e_/*N* values (Palstra and Ruzzante 2008; Clarke *et al*. 2024), which can be partly explained by either the big old fat fecund female fish hypothesis (Hixon *et al*. 2014) or the sweepstakes reproductive success hypothesis, which posits that only some families successfully reproduce (Hedgecock and Pudovkin 2011). A small *N*_e_/*N* implies a large variance in offspring number, leading to greater fluctuations in census size, which in turn threatens population persistence. However, there is currently no theoretical framework that explicitly connects *N*_e_ and *N*_e_/*N* to population persistence in the context of population viability analysis (PVA).

Since population persistence is inherently influenced by stochastic processes, understanding the role of stochasticity in population dynamics is crucial in PVA. Such stochasticity can be divided into two categories, apart from observation error: demographic and environmental stochasticity (Lande *et al*. 2003). Demographic stochasticity is due to individual-level events that occur within a reproductive period, such as births and deaths (Melbourne 2012). Environmental stochasticity is due to events that affect population-level or mean individual fitness and occur between reproductive periods, such as temperature changes, exploitation, density-dependent regulatory processes, competitive interactions, and other external factors (Ripa 2012). In sufficiently large populations, environmental stochasticity dominates population stochasticity, and demographic stochasticity can be neglected. However, in small populations, such as threatened species or subdivided populations, demographic stochasticity can affect population persistence even when the external environment is suitable for survival and reproduction, potentially leading to a greater chance of extinction. Given these risks, understanding how demographic and environmental stochasticity interact and contribute to overall population variability is crucial. Therefore, estimating the relative impact of these stochastic factors on population dynamics is essential in PVA (Lande 1993).

One indicator of how stochasticity affects population persistence is the magnitude of variance in recruitment to the next generation, which is shaped by both demographic and environmental factors (e.g. Lande 1993; Ovaskainen and Meerson 2010). Melbourne and Hastings (2008) theoretically derived the variance in recruitment by incorporating several stochastic factors. They showed that overlooked demographic factors, such as sex determination and fertility heterogeneity, significantly impacted the magnitude of the variance and, consequently, the stochastic extinction risk. Furthermore, they assessed the extinction risk for a laboratory population of semelparous species (i.e. a generation not overlapped) by incorporating these overlooked factors and estimating several parameters from experimental data. However, reliable estimates of demographic stochasticity parameters for wild populations are often difficult to obtain due to data limitations, particularly for threatened species. Additionally, demographic heterogeneity is expected to have a greater influence on iteroparous species (i.e. a generation overlapped by multiple reproductive cycles) than on semelparous species due to age structure (e.g. age-specific survival and/or fecundity), and collecting such data is even more challenging in the wild (Morris and Doak 2002).

For a population of iteroparous species, effective number of breeders (*N*_b_, Waples 1990) is a metric representing the contemporary effective size in a single breeding season, obtained by converting current *N* (hereafter referred to as the adult number), which is analogous to contemporary *N*_e_. *N*_b_ is defined based on known pedigree (Wang 2009; Waples and Waples 2011) or life-history information (Waples *et al*. 2011, 2013) and can be estimated from a single cohort sample using pedigree or life-history data or linkage disequilibrium (LD) information from genetic analyses (Waples 2006). Similar to *N*_e_/*N, N*_b_/*N* reflects the reproductive variance among parents (e.g. Waples 2002, 2024a). A small *N*_b_/*N* indicates over-dispersed reproduction caused by several factors. These factors can be partitioned into parental factors (e.g. age-/weight-specific fecundity) and non-parental factors (e.g. family-correlated survivorship after eggs hatching) (Akita 2020a,b), both of which are additional components of demographic stochasticity. Thus, it is reasonable to consider that *N*_b_/*N* provides information on demographic stochasticity and can be used to assess population persistence and extinction risk. It should be noted that, since most of the effects of life-history parameters and age-structure on population persistence are incorporated into *N*_b_, this methodological direction provides a potential approach for assessing population viability, especially for species with limited demographic data. However, there is little theoretical basis for using *N*_b_/*N* (or *N*_b_) to quantify demographic stochasticity.

In this study, we propose a theoretical framework for integrating *N*_b_ and *N*_b_/*N* into the stochastic dynamics of the number of adults (*N*) for iteroparous species, including both demographic and environmental stochasticity. To achieve this, we derive a theoretical expression for recruitment process by decomposing it into three stochastic components: i) parental components, which influence reproductive output; ii) non-parental components, which affect early survival through family-correlated effects; and iii) environmental components, which shape overall survival. This approach allows us to assess the relative contributions of demographic and environmental stochasticity, with *N*_b_ summarizing the effects of the first two components (Akita 2020a).

Our modeling framework can be applied to diverse diploid species under an arbitrary mating system regardless of semelparity or iteroparity. For semelparous species, a change from *N*_b_ to *N*_e_ ensures consistency in the results. However, the description of the model focuses on iteroparous fish species, which are among the best candidates for applying the proposed method.

## Theory

Here, we present the theoretical basis for integrating *N*_b_ into stochastic population dynamics. The main contribution of this study to the existing literature is the derivation of an equation for recruitment variation (Equations 10–11). This set of equations forms the basis for a new index of demographic stochasticity, which quantifies the degree to which it is amplified by environmental stochasticity. Details of the derivation are provided in the **Appendix**; this section focuses on the model settings and assumptions. The main symbols used in this section are summarized in Table 1.

**Table 1.**
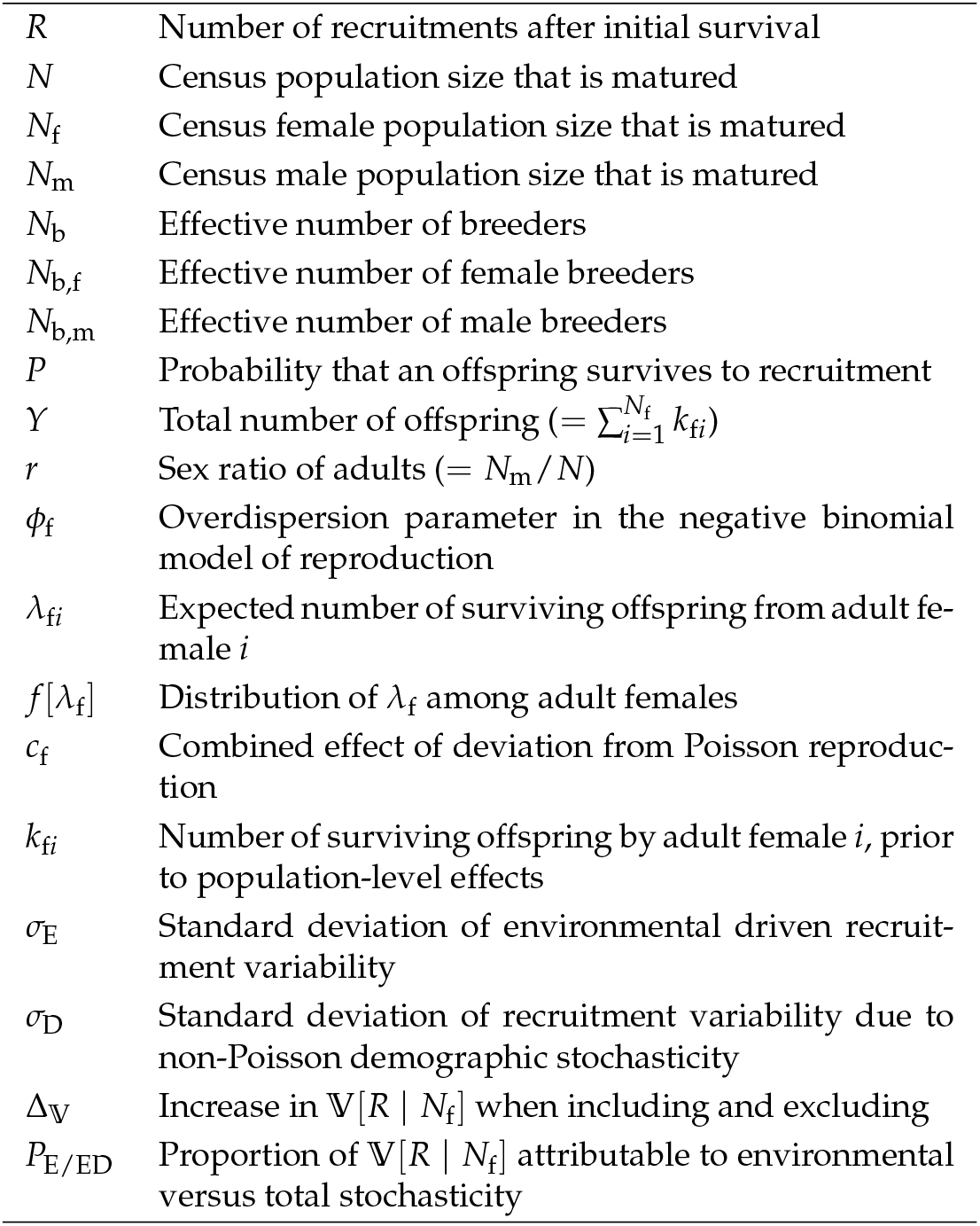
List of mathematical symbols used in the main text.

### Overview

We consider a hypothetical population of semelparous species with *N* parents, assuming no population subdivision or spatial structure. A mature individual is referred to as a parent or adult, even if it does not produce offspring. The total number of parents is denoted as the census size, following standard usage in conservation biology. We examine the stochastic process by which the recruitment number (*R*) is generated from *N* parents during a single reproductive season.

A common equation in fisheries science (Hilborn and Walters 1992) describes this relationship as

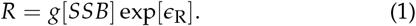

where *SSB* is the spawning stock biomass (= ∑_*a*_ *N*_*a*_*W*_*a*_, with *N*_*a*_ and *W*_*a*_ representing the number and weight of parents at age *a*, respectively). The function *g*[] represents the stock– recruitment relationship, including density dependence, such as in the Beverton–Holt or Ricker models. The term *ϵ*_R_ represents Gaussian noise with mean 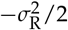 and variance 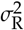, such that the expectation of *R* is *g*[*SSB*] (denoted by 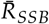). The variance of *R* is given by

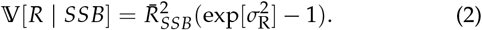

If 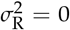, then 𝕍[*R* | *SSB*] = 0, indicating that the recruitment process is deterministic.

Here, 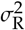 summarizes population-level events, such as environmental fluctuations, that affect all individuals similarly, and is therefore interpreted as environmental stochasticity. However, this top-down approach, based solely on population-level dynamics (Equation 1), does not incorporate demographic stochasticity, which arises from individual-level events.

To address this limitation, we develop a bottom-up approach based on family-level processes linking adult females, eggs, offspring, and recruitment. Among the *N* parents, let *N*_f_ be the number of adult females before spawning. Each adult female produces eggs according to her reproductive potential *λ*_f_, which varies across adult females and follows a distribution *f* [*λ*_f_] (subscript “f” denotes female). Using this model, we derive the expression for 𝕍[*R* | *N*_f_, *f* [*λ*_f_]] that incorporates both demographic and environmental stochasticity. For simplicity, we hereafter denote this as 𝕍[*R* | *N*_f_]. It is worth noting that 𝕍[*R* | *N*_f_, *f* (*λ*_f_)] reduces to 𝕍[*R* | *SSB*] if we ignore demographic stochasticity and the male contribution to *SSB*, and assume that *f* (*λ*_f_) is the distribution of female weights, i.e., *SSB* = *N* ∫ *λ*_f_ *f* (*λ*_f_)*dλ*_f_.

We further examine how the effective number of female breeders, denoted by *N*_b,f_, affects 𝕍[*R* | *N*_f_]. Assuming an arbitrary degree of environmental stochasticity, we theoretically demonstrate that small values of *N*_b,f_ and *N*_b,f_/*N*_f_ (or *N*_b_ and *N*_b_/*N*) amplify the contribution of environmental stochasticity to recruitment variance.

### Bottom-up approach

To incorporate individual-level events into recruitment variation, we propose a bottom-up approach based on an individual-based model, following previous studies (e.g. Engen *et al*. 1998; Brännström and Sumpter 2006; Melbourne and Hastings 2008; Kendall and Wittmann 2010; Ferguson and Ponciano 2014, 2015). The model is designed to capture the stochastic processes underlying reproduction, early-stage survival, and late-stage survival leading to recruitment.

The model consists of three stochastic processes. The first is the parental process, in which parental individuals produce eggs according to their reproductive potential (e.g. age or size). The second is the non-parental process, where early-stage survival is influenced by parental differences, such as family-correlated survival after hatching, even when reproductive potential is similar among parents. The third is the environmental process, which governs late-stage survival under equal survival probabilities among offspring in the absence of parental or familial influence. This environmental process is driven by population-level events, including density-dependent effects, commonly described by survival functions such as the Beverton–Holt or Ricker model. Figure 1 illustrates a schematic representation of the recruitment process in fish, showing how the parental, non-parental, and environmental processes correspond to reproduction, early-stage survival, and late-stage survival, respectively. Here, we define the parental and non-parental processes as demographic processes.

**Figure 1.**
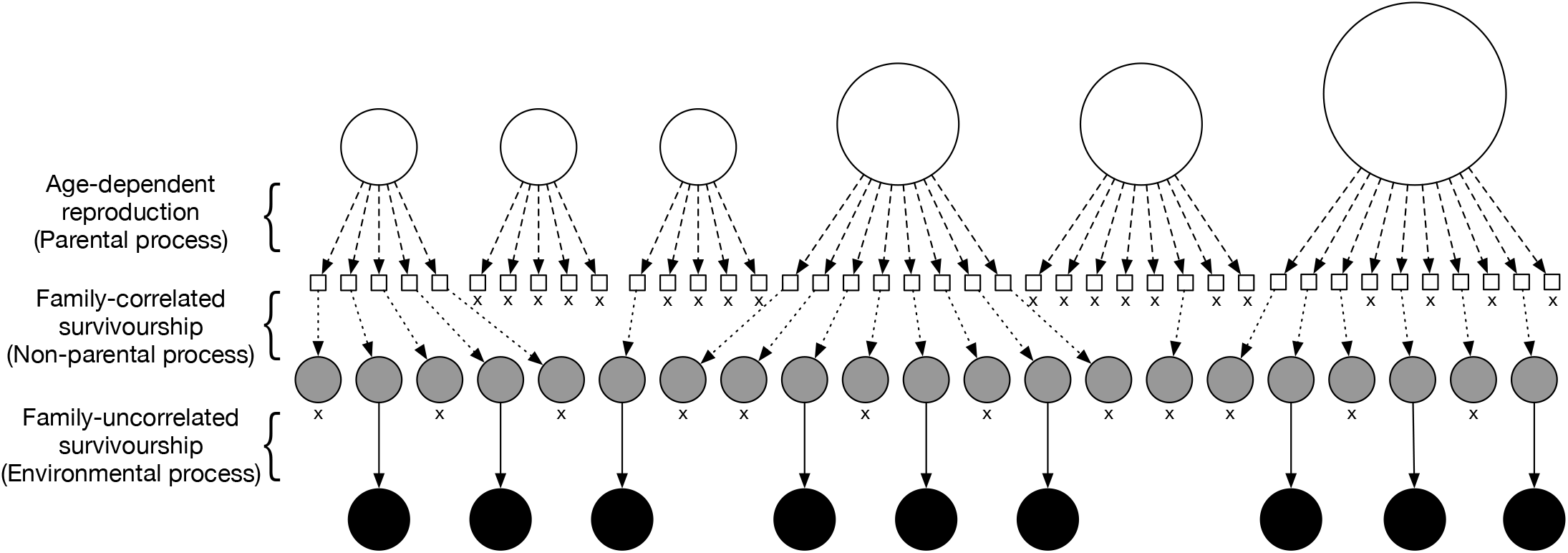
Relationships among adult females, eggs, offspring and recruitments. The open circles, open squares, gray circles, and filled circles represent adult females, eggs, offspring, and recruitment respectively. The area of an open circle indicates the degree of reproductive potential of an adult female (i.e. *λ*_f*i*_). The dashed, dotted, and thin arrows indicate parent–egg, egg–offspring, and offspring–recruitment relationships, respectively. The symbol “× “indicates failure to survive till the recruitment stage.

### Reproductive variance governed by parental and non-parental factors

We examined how demographic processes affect the variance in offspring number per parent, focusing on the maternal lineage. For simplicity, we considered only adult females and did not distinguish the sex of offspring. Let *k*_f*i*_ denote the number of offspring that were produced by adult female *i* (*i* = 1, 2, …, *N*_f_) and have passed the demographic processes but not yet entered the late-stage survival phase (gray circles in Figure 1). Following Akita (2020a,b), *k*_f*i*_ was assumed to follow a negative binomial distribution with a mean of *λ*_f*i*_ :

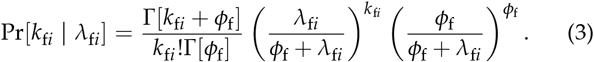

Here, *λ*_f*i*_ is independently drawn from a distribution *f* [*λ*_f_] that captures parental variation. *λ*_f*i*_ represents the reproductive potential of each female, which may reflect physiological or behavioral states related to reproductive success, such as maternal size, weight, accumulated energy reserves, hatching days, or residence time on the spawning ground. The overdispersion parameter *ϕ*_f_ describes non-parental sources of variation (e.g. family-correlated survival). For mathematical tractability, *ϕ*_f_ is assumed to be constant across adult females (i.e. parental and non-parental factors are determined independently).

Akita (2020a,b) derived the variance in the number of surviving offspring per adult female as

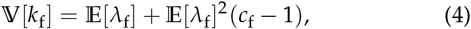

where

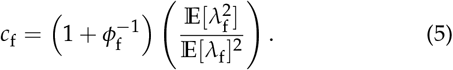

The coefficient *c*_f_ reflects the combined effects of parental and non-parental variation, and it satisfies *c*_f_ ≥ 1. When either source of variation is absent (i.e. *c*_f_ → 1), 𝕍[*k*_f_] reduces to 𝔼[*λ*_f_], which corresponds to Poisson variance in female reproductive output.

Following Akita (2020a), we define *N*_b,f_ as the inverse of the probability *π*_f_ that two offspring share a maternal sibling relationship with any adult female:

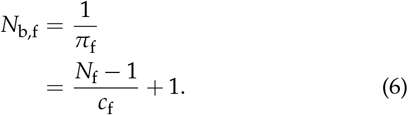

In this formulation, *N*_b,f_ is independent of the mean reproductive potential 𝔼[*λ*_f_], but it depends on *ϕ*_f_ and the coefficient of variation ℂ𝕍[*λ*_f_], since 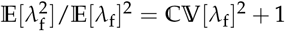.

Notably, this formulation of *N*_b,f_ is based on maternal sibship probabilities and is conceptually related to the well-known estimator derived by Wang (2009), which considers contributions from both sexes to estimate the overall *N*_b_. In contrast, our approach focuses exclusively on one sex (female) and allows the variance in reproductive success to be explicitly partitioned into parental and non-parental components. This decomposition enhances the interpretability of the demographic contributions to *N*_b,f_ and provides a clearer link between biological processes and stochastic variation in recruitment.

Furthermore, following Akita (2020b), we approximate the ratio of *N*_b,f_ to *N*_f_ as

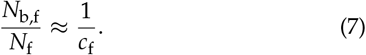

When either source of variation exists (i.e. *c*_f_ > 1), *N*_b,f_/*N*_f_ is less than one. Conversely, in the absence of both sources of variation (i.e. under Poisson reproduction), *N*_b,f_/*N*_f_ equals unity. Thus, *N*_b,f_/*N*_f_ reflects the degree of deviation from Poisson-like reproduction. In our model, we assume *N*_f_ ≥ *N*_b,f_ ≥ 2.

### Recruitment variation driven by environmental stochasticity

We next describe how environmental stochasticity contributes to recruitment variation (see thin arrows in Figure 1). After initial survival governed by demographic processes, we assume that each offspring has an equal probability of surviving to recruitment, in the absence of parental or family-level influence. The recruitment number *R* is modeled as a binomially distributed random variable:

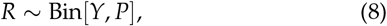

where *Y* is the total number of offspring 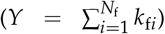 and *P* is the density-dependent survival probability. We assume that *Y* and *P* are independent, since the model separates the population-level density-dependent phase from the earlier, individual-driven survival phase. We define this phase as the late-life stage.

To incorporate environmental stochasticity, we model *P* as a log-normal random variable:

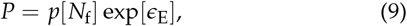

where *ϵ*_E_ is a normally distributed random variable with mean 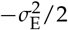 and variance 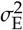. This ensures that 𝔼[*P* | *N*_f_] = *p*[*N*_f_], where *p*[*N*_f_] represents the expected density-dependent survival. For simplicity, we hereafter denote 𝔼[*P* | *N*_f_], 𝔼[*Y* | *N*_f_], and 𝔼[*R* | *N*_f_] as 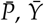 and 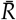 respectively, where 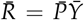 follows from the assumed independence of *Y* and *P*.

### Model flexibility and limitations

While our modeling framework is designed to flexibly capture a wide range of reproductive ecological phenomena, it also involves several assumptions and limitations. Below, we outline these considerations, focusing particularly on the interpretation and application of *N*_b,f_.

First, in our model based on the inbreeding rate (Akita 2020a), *N*_b,f_ is determined by the number of offspring per parent observed after surviving the early-life stages, incorporating variation in reproductive potential and family-correlated survival. In other words, *N*_b,f_ remains theoretically consistent during the late-life stage and can be estimated regardless of the timing of offspring sampling, as long as all surviving offspring are sampled with equal probability during this period. This consistency is a practical advantage. That said, the assumption of independence among the three sources of variation may not always hold, for example, if larger females are more likely to access rare, high-quality spanwing grounds, or if the effects of reproductive potential or family-correlated survival persist beyond the early-life stages and influence recruitment.

In contrast, the effective number of breeders based on allele frequency variance is more sensitive to sampling timing, particularly in species with type III survivorship curves. In such species, overdispersed offspring production during early-life stages can inflate the variance-to-mean ratio, complicating the interpretation of effective size unless the variance is appropriately rescaled (Crow and Morton 1955; Waples 2002).

Second, although our model assumes no explicit spatial structure or dispersal, it can accommodate fine-scale reproductive variation that influences reproductive success or early survival by treating such effects as part of the non-parental component. For example, local environmental heterogeneity at spanwing grounds, such as that shaped by maternal habitat choice, can be absorbed into the family-correlated survival term. The phenomenon of skipped spawning, commonly observed in iteroparous fishes, is also implicitly captured through variation in reproductive potential: adult females that do not reproduce in a given season are effectively represented as having near-zero *λ*_f_. Because *N*_b,f_ is defined per reproductive season, such behavioral variation is naturally reflected in seasonal estimates. This also implies that the estimate may not represent the entire adult female population if some individuals do not reproduce during the focal season (Waples and Feutry 2022). The same framework is equally applicable to male breeders. If energetic investment in reproduction differs between sexes, especially across years, such asymmetry can be reflected in the difference between female and male effective numbers of breeders.

Finally, our definition of *N*_b_ can be used as an indicator of inbreeding depression, but does not explicitly include inbreeding itself. This limitation should be kept in mind when interpreting the theoretical results presented later. In contrast, the formulation of *N*_b_ derived by Wang (2009) incorporates deviations from Hardy–Weinberg proportions and thus explicitly captures the effects of inbreeding. However, because *N*_b_ is constructed from more limited information than *N*_e_, it may not allow accurate quantification of the full effects of inbreeding (e.g. Waples and Antao 2014; Whiteley *et al*. 2015). In any case, when applying our definition of *N*_b,f_ (and *N*_b_) to small populations, it is important to recognize that ignoring inbreeding could lead to upward bias.

## Results

### Partitioning of recruitment variations

Given *N*_f_ and the functional relationship between *N*_f_ and 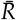, the variance in recruitment can be expressed as the combined effects of demographic and environmental processes:

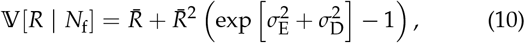

where

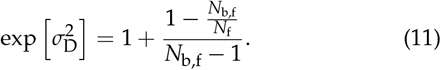

The interpretation of 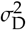 is discussed below, and the full derivation of Equations 10–11 is provided in **Appendix**. This variance expression implies that, given *N*_f_, *R* follows a Poisson lognormal distribution with mean 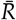.

When 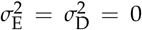, the variance reduces to 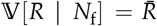, corresponding to the Poisson variance in recruitment. By contrast, when 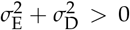, most of the variance arises from the second term in Equation 10. The components 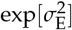 and 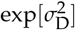 act synergistically to amplify the variance, even in the absence of environmental variation (i.e. when 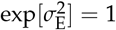).

Equation 11 shows that 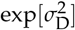 decreases monotonically as *N*_b,f_ increases: when 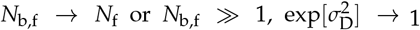, indicating that the recruitment variance is governed only by Poisson and environmental factors. When *N*_b,f_ < *N*_f_, it reflects a non-Poisson reproductive process influenced by parental and/or non-parental variation. Therefore, 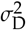 can be interpreted as a non-Poisson demographic component. However, its contribution may diminish with increasing *N*_b,f_ even when *N*_b,f_ < *N*_f_, as reflected in Equation 11, where the large denominator of the second term reduces the influence of non-Poisson reproductive variation, indicating a reduction in finite size effects.

It is worth noting that Equation 10 reduces to the form of the top-down model (Equation 2) when the demographic components are excluded, as expected. This agreement confirms that the proposed bottom-up model generalizes the standard population-level formulation by incorporating individual-level stochasticity.

In summary, the recruitment variance in Equation 10 can be partitioned into (i) a Poisson demographic component (the first term) and (ii) a combined component reflecting non-Poisson demographic and environmental stochasticity (the second term). The influence of the non-Poisson demographic component becomes especially pronounced when both *N*_b,f_/*N*_f_ and *N*_b,f_ are small.

When the second term dominates, the coefficient of variation (CV) of recruitment can be approximated as

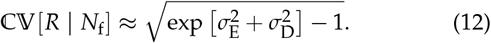

In this formulation, the dependence on 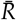 is omitted, highlighting that the CV is approximately independent of the specific functional form of the relationship between *N*_f_ and 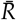, including density dependence.

### Application to species with low *N*_b_/*N* ratios

As demonstrated in the previous subsection, our model predicts that the contribution of non-Poisson demographic variation to recruitment variance becomes non-negligible when both *N*_b_/*N* and *N*_b_ are small. Recent meta-analyses (Clarke *et al*. 2024) have identified taxa and species with notably low *N*_b_/*N* ratios, raising the possibility that our model could be used to quantitatively assess how reductions in *N*_b_ impact recruitment variance in these groups. However, because our current formulation is female-specific, direct comparisons to empirical *N*_b_ values, which typically include contributions from both sexes, are not straightforward. To bridge this gap, we introduce a simplified extension of our model to include both sexes under a set of coarse assumptions. This allows us to reformulate the model in terms of *N*_b_ and *N*, and propose a rough quantitative guideline for interpreting the potential influence of non-Poisson reproduction in species with low *N*_b_/*N* ratios.

### Extending the model to both sexes under simplifying assumptions

The effective number of breeders *N*_b_ can be derived by combining the effective number of female breeders (*N*_b,f_) and that of male breeders, denoted by *N*_b,m_, following the standard formulation (Crow and Denniston 1988; Caballero 1994):

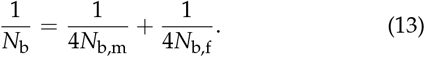

Assuming an equal adult sex ratio, such that *N*_f_ = *N*_m_ = *N*/2, and no sex differences in reproductive variation (i.e., *N*_b,m_ = *N*_b,f_), Equation 13 reduces to *N*_b,f_ = *N*_b_/2. We can substitute these into Equation 11 to obtain a reformulated expression:

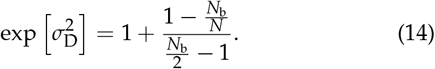

This expression holds under the following assumptions: (i) the probabilities that two sampled offspring share a mother or a father are equal, and (ii) the adult sex ratio is exactly 0.5.

This equation is particularly useful for estimating 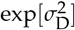 when maternal and paternal sibships cannot be reliably distinguished, allowing *N*_b_ to be estimated without separately inferring *N*_b,f_ and *N*_b,m_. This feature is advantageous because distinguishing between maternal and paternal sibships can be difficult and may introduce additional uncertainty. Note that this formulation assumes *N* ≥ *N*_b_ ≥ 4.

### *N*_b_ as an indicator of non-Poisson amplification of recruitment variance

As demonstrated in the previous subsection, the non-Poisson demographic factor 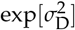 can be expressed as a function of *N*_b_ and *N* under simplifying assumptions (Equation 14). Figure 2a illustrates this relationship for different values of *N*. As *N*_b_ increases, 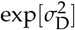 decreases and asymptotically approaches unity, indicating that the contribution of non-Poisson effects to recruitment variance becomes negligible when *N*_b_/*N* is close to one or *N*_b_ exceeds 100.

**Figure 2.**
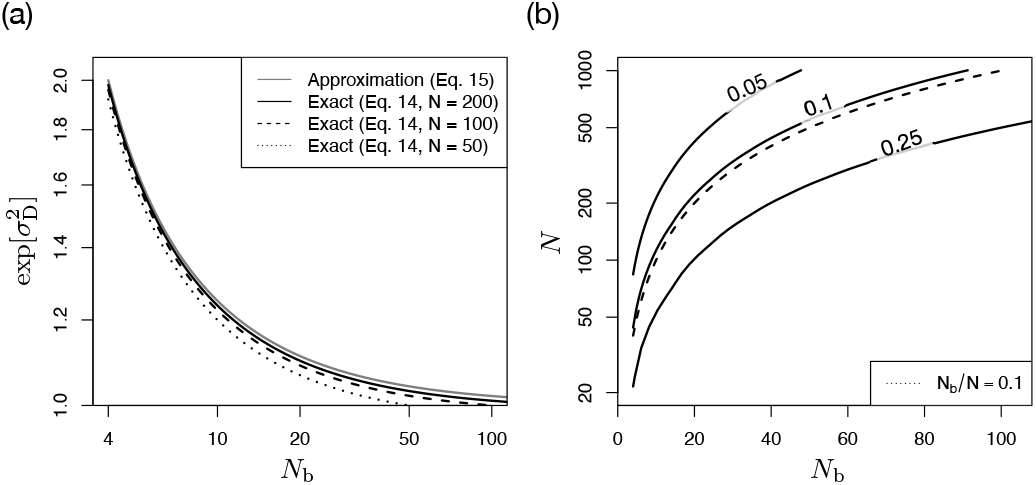
(a) Relationship between the effective number of breeders (*N*_b_) and the non-Poisson demographic factor 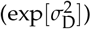. The bold, dashed, and dotted lines indicate the exact relationship in Equation 14 at *N* =200, 100, and 50 respectively. The gray line indicates the approximated relationship in Equation 15. The x- and y-axes are presented in the logarithmic scale. (b) Contour plot of the approximation accuracy in Equation 15 as a function of *N*_b_ and *N*. The contour labels in the plot indicate the value of a relative approximation error, which is defined as (approximated value exact − value)/exact value. The dotted line represents the relationship that satisfies *N*_b_/*N* = 0.1. The y-axis is presented in the logarithmic scale.

**Figure 3.**
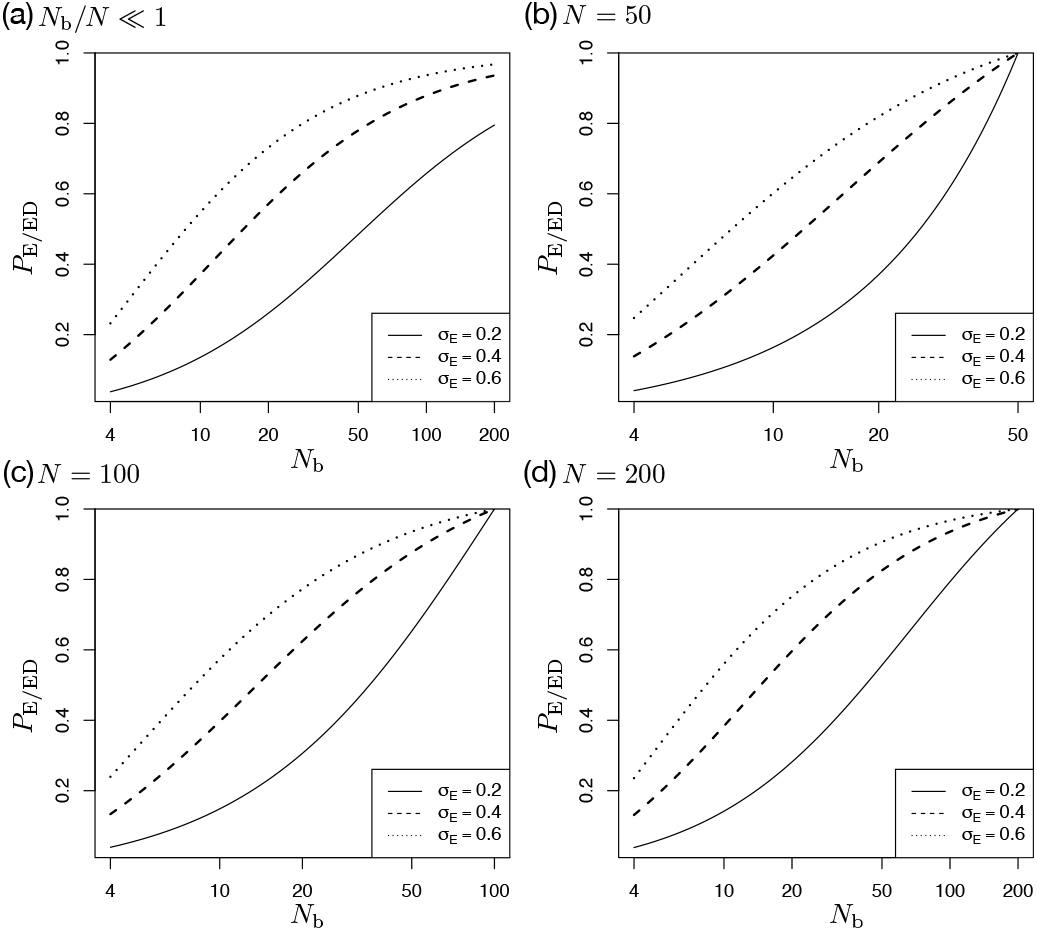
Relationship between the effective number of breeders (*N*_b_) and the ratio indicating the extent to which the variance in recruitment is underestimated when non-Poisson effects are ignored (*P*_E/ED_). The bold, dashed and dotted lines correspond to *σ*_E_ = 0.2, 0.4, and 0.6 respectively. The x-axis is presented in the logarithmic scale. (a) Approximate relationship using Equation 15 (*N* is not specified). (b-d) Exact relationship using Equation 14 (*N* is specified).

Conversely, when *N*_b_ is below approximately 50 and the *N*_b_/*N* ratio is small, 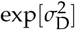 remains appreciably greater than one, and the influence of the non-Poisson factor cannot be ignored.

If *N*_b_/*N* is sufficiently small, 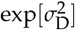 can be approximated by:

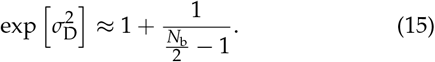

Figure 2b shows the contour plot of the relative approximation error. As expected, the error decreases as *N*_b_/*N* becomes smaller. In particular, the region above the dotted line (*N*_b_/*N* < 0.1) closely corresponds to areas where the approximation error is below 10%, indicating that the approximation in Equation 15 is likely acceptable for practical use. In such cases, the value of *N* is not required to estimate 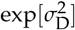.

Therefore, if *N*_b_ is available and falls within this range, Equation 15 offers a convenient means to quantify the influence of non-Poisson demographic stochasticity (gray line in Figure 2a). These results highlight the utility of *N*_b_ as a sufficient metric for evaluating the contribution of demographic stochasticity to recruitment variance beyond Poisson expectations, especially in scenarios where the influence of *N* can be ignored. It is important to note that this approximation consistently yields upwardbiased estimates of 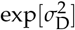, and thus may slightly overestimate the contribution of demographic stochasticity to recruitment variance.

To further illustrate how non-Poisson demographic variation contributes to recruitment variance, we evaluated its impact from two complementary perspectives. First, we quantified the increase in recruitment variance attributable to the non-Poisson component by comparing scenarios with and without this factor.

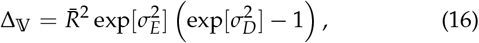

This expression shows that Δ_𝕍_ increases in proportion to 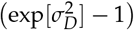, as 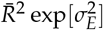 and 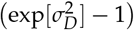 are independently determined given *N*. For example, using the approximation in Equation 15, when *N*_b_ = 4, 8, 22, and 42, Δ_𝕍_ increases by approximately 100%, 33%, 10%, and 5%, respectively (gray line in Figure 2a).

Second, we evaluated the extent to which the variance in recruitment is underestimated when the non-Poisson component is ignored. To quantify this, we introduced the ratio *P*_E/ED_:

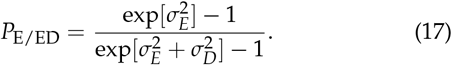

This ratio compares the contribution of environmental stochasticity alone (numerator) with the combined contribution of environmental and demographic stochasticity (denominator) to the total recruitment variance. A value close to unity indicates that the demographic component adds little to the overall variation, while lower values indicate a substantial contribution of non-Poisson demographic stochasticity. Note that This formulation assumes 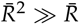.

Figure 17a shows the relationship between *N*_b_ and *P*_E/ED_ for different levels of environmental stochasticity (*σ*_*E*_ = 0.2, 0.4, 0.6), using the approximation in Equation 15. As expected, *P*_E/ED_ approaches unity as *N*_b_ increases, and this convergence is more rapid under stronger environmental stochasticity. Conversely, when *N*_b_ is small, the demographic component has a larger impact. For example, at *N*_b_ = 50 and *σ*_*E*_ = 0.2, *P*_E/ED_ = 0.48, indicating that more than half of the variance is attributable to demographic stochasticity. In comparison, when *σ*_*E*_ = 0.6, *P*_E/ED_ = 0.88, suggesting that the demographic effect is weaker but still non-negligible. These patterns were derived under the assumption that *N*_b_/*N* is sufficiently small (e.g. < 0.1). However, when both *N*_b_ and *N* are known, Equation 14 allows for direct computation without relying on this approximation (Figure 17b–d, for *N* = 50, 100, and 200, respectively).

Furthermore, when the aforementioned approximation is valid, replacing 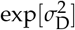 in Equation 12 shows that the coefficient of variation (CV) depends primarily on *N*_b_ and 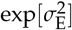.

## Discussion

This study introduces a conceptual shift in how the effective number of breeders (*N*_b_) is interpreted within ecological and evolutionary frameworks. Traditionally treated as a genetic summary statistic derived from pedigree or genotype data, *N*_b_ has not typically been embedded within the explicit mechanics of demographic stochasticity. Here, we present a mechanistic model in which *N*_b_, specifically its female component *N*_b,f_, is formulated as a function of reproductive variance arising from parental and non-parental processes, which together constitute the additional demographic component of recruitment variability. By making this connection explicit, our framework unifies two concepts that have historically evolved in parallel: effective population size (defined through reproductive variance and central to evolutionary genetics), and demographic stochasticity (which underpins population viability theory). Thus, *N*_b_ or *N*_b,f_ provides a direct measure of how strongly demographic stochasticity, arising from non-Poisson reproductive variation, can amplify recruitment variance, particularly in small or declining populations.

To support this conceptual shift, we provided a theoretical basis for integrating *N*_b_ into stochastic population dynamics for iteroparous species, grounded in individual-level relationships between adult females and their offspring. Recruitment variation was analytically derived by partitioning the recruitment process into three components: parental, non-parental, and environmental. The parental and non-parental components jointly define the demographic stochasticity that arises from individual heterogeneity. The resulting expression for recruitment variance (Equation 10) consists of a term proportional to 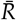, reflecting Poisson demographic variance, and a second term proportional to 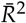, which captures the synergistic amplification due to non-Poisson reproductive variation and environmental stochasticity. This second term dominates recruitment variance in most realistic scenarios. Assuming a 1:1 sex ratio and no sex differences of reproductive variance, we derived a general relationship between *N*_b_ and 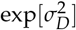 (Equation 14), and provided a simplified approximation under the condition that *N*_b_/*N* ≪ 1 (Equation 15). This theoretical framework enables practical interpretation of recruitment variability in relation to *N*_b_, with or without knowledge of total census size.

In the following, we treat *N*_b_ as a practical proxy for contemporary *N*_e_, consistent with its common use in conservation genetics.

### Bridging demographic viability and genetic health

The causal relationship between genetic diversity and population viability has long been debated (Kardos *et al*. 2021; Teixeira and Huber 2021), yet empirical evidence for a strong link remains limited. Several genetic mechanisms have been proposed to explain population decline, including inbreeding depression (e.g. Hedrick and Garcia-Dorado 2016), mutation load (e.g. Agrawal and Whitlock 2012), and maladaptation to changing environments (e.g. Brady *et al*. 2019). These models typically operate through reductions in individual fitness, which in turn lead to lower survival or reproductive success at the population level.

In contrast, our framework highlights a distinct causal pathway, which is relevant to ongoing discussions about whether losses of genetic diversity are drivers or consequences of population decline (Lande 1995; Frankham 2005). Our model analytically demonstrates that demographic stochasticity can be amplified by reproductive variance exceeding Poisson expectations, and that this amplification is directly measurable through the effective number of breeders, *N*_b_, if *N*_b_/*N* is small. This amplification increases recruitment variance and, in turn, may reduce population persistence, providing a novel theoretical perspective that supports the interpretation that losses of genetic diversity, resulting from increased reproductive variance, can act as a driver of population decline. Notably, the form of *N*_b_ used in this study provides a neutral proxy for this demographic component, as it is derived solely from individual-level reproductive variance and does not incorporate fitness-related effects.

This perspective complements recent efforts to integrate genetic and demographic indicators in conservation biology (Shaw *et al*. 2025). Our model highlights a conceptual expansion of *N*_b_: beyond its role as a neutral genetic statistic, it also serves as a quantitative indicator of demographic risk. This dual role positions *N*_b_ as a bridge between genetic monitoring and population viability analysis (PVA).

### Scope and limitations for species with low *N*_b_/*N* ratios

Species with exceptionally low ratios of effective to census population size (*N*_b_/*N*), such as many marine organisms and insects, are particularly relevant to the framework we present. Recent meta-analyse has shown that in these taxa, *N*_b_/*N* often falls below 0.1 (Clarke *et al*. 2024), placing them within the parameter space where our theory predicts that recruitment variance may be strongly amplified by the interaction of non-Poisson demographic factors and environmental stochasticity.

Our results indicate that when *N*_b_ is small, on the order of 50 or fewer, the amplification of recruitment variance becomes substantial. Clarke *et al*. (2024) reported that the groups with the lowest mean *N*_b_ estimates were diadromous fishes (*N*_b_ = 33), amphibians (*N*_b_ = 47), and reptiles (*N*_b_ = 49), suggesting that our framework is applicable to many of these species, provided that *N*_b_/*N* < 0.1 also holds.

It is important to note, however, that many species with low *N*_b_/*N* ratios, especially marine fishes, also exhibit large values of *N*_b_, often exceeding the critical threshold (e.g. *N*_b_ < 100) (Clarke *et al*. 2024), above which demographic stochasticity contributes minimally to recruitment variance. For these species, the demographic risks we highlight may not be immediately relevant under typical natural conditions.

Nonetheless, populations of such species may become vulnerable when fragmented into small subpopulations or newly established through colonization events. In these scenarios, *N*_b_ can fall below the critical range, and our framework suggests that demographic stochasticity, alongside known genetic threats such as inbreeding depression or mutational load, may further reduce population viability. This insight offers a new perspective for evaluating persistence in spatially structured populations, including metapopulations and reintroduction scenarios.

Moreover, for species inhabiting terrestrial or freshwater environments, where overall census sizes are typically more moderate, species with low *N*_b_/*N* ratios may fall directly within the vulnerable zone (Clarke *et al*. 2024). In such contexts, non-Poisson demographic stochasticity should be considered alongside other recognized extinction risk factors, particularly in populations experiencing environmental instability or anthropogenic stress.

### Life-history-independent assessment of demographic stochasticity

One of the key strengths of our framework is that it provides a quantitative basis for assessing the contribution of demographic stochasticity to recruitment variation across species, regardless of their life-history traits. Traditional approaches to estimating demographic variance typically require detailed demographic or reproductive data, which vary widely among taxa and often complicate cross-species comparisons. In contrast, our results show that the magnitude of demographic stochasticity driven by overdispersed reproduction can be inferred directly from the effective number of breeders, *N*_b_, particularly when *N*_b_/*N* is small. This offers a unified and lifehistory-independent basis for comparing vulnerability across species with vastly different reproductive schedules and lifespans. While the consequences of a given level of recruitment variance for extinction risk may still depend on species-specific traits such as generation time and recovery potential (Provost and Botsford 2022), our framework isolates a core demographic component that is broadly comparable across species.

This interpretation has particular relevance in the context of PVA, where extinction risk is often assessed based on the time required for a population to decline to a quasi-extinction threshold (e.g. Holmes *et al*. 2007). A common practice in PVA is to define this threshold at a population size below which demographic stochasticity becomes a dominant force in extinction dynamics (Morris and Doak 2002). However, the critical size at which this shift occurs can vary substantially among species due to differences in life-history traits (e.g. Fujiwara 2007), making it difficult to establish general criteria across taxa. By linking this demographic sensitivity directly to *N*_b_, our framework offers a universal reference point for identifying the population sizes at which demographic stochasticity is likely to become consequential, without requiring life-history-specific parameterization.

### Operational utility and caveats in *N*_b_ estimation

A practical advantage of using *N*_b_ for this purpose is that it can be estimated from genetic data alone, without requiring information on life-history parameters or the current age/size structure of the population. Specifically, *N*_b_ can be inferred from samples drawn from a single cohort. This feature is especially useful in conservation contexts, where endangered species often lack comprehensive demographic data and sampling adult individuals may be difficult or constrained. For such species, genetic samples from a single cohort of offspring may suffice to estimate *N*_b_ using sibship frequency methods (Wang 2009) or linkage disequilibrium (LD) methods (Waples 2006). These methods yield estimates of *N*_b_ for either the parental generation or, in the case of LD-based approaches, for an average over recent generations (Wang *et al*. 2016).

While estimating *N*_b_ in very large populations is inherently difficult due to weak signals of drift or inbreeding (Wang 2025; Waples 2024b), our framework is most relevant to populations where *N*_b_ is small (e.g. below a few hundred). In these cases, *N*_b_ estimates are more precise and interpretable, and they directly inform the expected degree of demographic stochasticity in recruitment due to non-Poisson component. A caveat is that when sampling very young individuals (e.g. young-of-year), biased sampling, family-correlated sampling (Waples 2025), could lead to inflated estimates of reproductive variance. It is therefore important to ensure sufficiently random and representative sampling of the cohort to avoid artificially inflating the inferred reproductive variance.

## Data availability

Code used to draw the figures can be found at https://github.com/teTUNAakita/DemoStoNb.

## Acknowledgments

The author is very grateful to H. Okamura and R.S. Waples for insightful comments on an earlier version of this manuscript. The author also thanks M. Ichinokawa for fruitful discussions.

## Appendix

### Derivation of 𝕍[*R* | *N*_f_] (Equations 10–11)

Suppose for iteroparous species there are *N*_f_ adult females in a closed population and heterogeneity in reproductive potential among them is allowed. Adult female *i* (*i* = 1, 2, …, *N*_f_) produces *k*_f*i*_ offspring that have experienced the initial survival events as a function of family-correlated effects. Of the total number of offspring, *Y*(= ∑ *k*_f*i*_), *R* individuals survive as recruitment with probability *P* through population-level events that are independent of family influence. Herein, we describe the derivation of the recruitment variation given the number of adult females, 𝕍[*R* | *N*_f_]. This is performed by computing the expectation, variance and second moment conditional on *N*_f_ for *Y* and *P*.

Following the distribution of *k*_f*i*_ | *λ*_f*i*_ (Equation 3 in the main text), the expectation of *Y* | *N*_f_ is given by

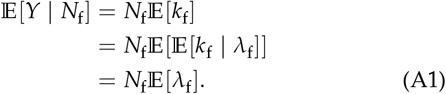

When computing the expectation or variance, we drop the subscript (“*i*”) because random variables, such as *k*_f_ and *λ*_f_, are independent and identically distributed. The variance and second moment for *Y* | *N*_f_ is given by

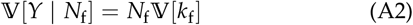

and

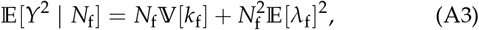

respectively. Following Akita (2020a,b), we can obtain 𝕍[*k*_f_] as

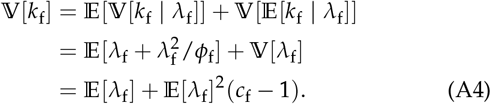

where *c*_f_ indicates the combined effect of deviation from the Poisson (Akita 2020a), defined by Equation 5 in the main text.

The survival probability of offspring to recruitment, *P*, is dependent on population-level effects and is assumed to be a random variable with a log-normal distribution.

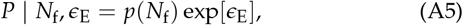

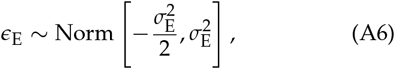

where *p*(*N*_f_) and *ϵ*_E_ correspond to the deterministic part of the density-dependent survival probability and stochastic noise, respectively. This equation provides a relatively simple expectation due to the bias correction treatment in Equation A6, such as

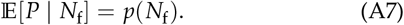

The variance and second moment for *P* | *N*_f_ is given by

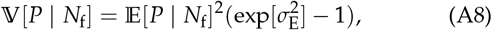

and

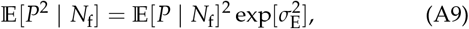

respectively.

The recruitment number, *R*, is assumed to be determined by a binomial process with the total offspring number (*Y*) and *P*.

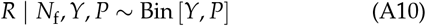

Given *N*_f_, it is natural to consider that *Y* and *P* are independently determined; thus, we can easily obtain the conditional expectations of *R* such as:

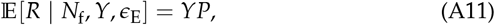

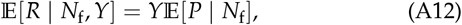

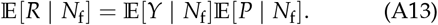

The conditional expectations of *R* required slightly more complex equations, such as

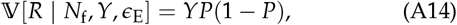

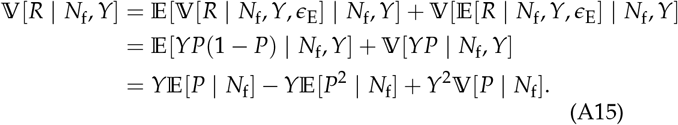

Finally, we obtain the recruitment variation given the number of adult females, such as

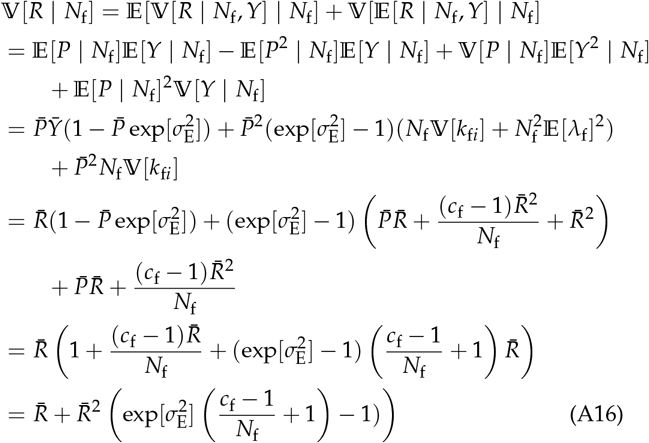

For clarity, we replaced 𝔼[*P* | *N*_f_], 𝔼[*Y* | *N*_f_] and 𝔼[*R* | *N*_f_] by 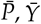, and 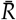(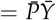, Equation A13), respectively. Using the relationship of *N*_b,f_ (Equation 6), we can obtain the Equations 10–11 i the main text.

